# An image segmentation method based on the spatial correlation coefficient of Local Moran’s I - identification of A-type potassium channel clusters in the thalamus

**DOI:** 10.1101/2023.05.02.539063

**Authors:** Csaba Dávid, Kristóf Giber, Katalin Kerti-Szigeti, Mihaly Kollo, Zoltán Nusser, László Acsády

## Abstract

Unsupervised segmentation in biological and non-biological images is only partially resolved. Segmentation either requires arbitrary thresholds or large teaching datasets. Here we propose a spatial autocorrelation method based on Local Moran’s I coefficient to differentiate signal, background and noise in any type of image. The method, originally described for geoinformatics, does not require a predefined intensity threshold or teaching algorithm for image segmentation and allows quantitative comparison of samples obtained in different conditions. It utilizes relative intensity as well as spatial information of neighboring elements to select spatially contiguous groups of pixels. We demonstrate that Moran’s method outperforms threshold-based method (TBM) in both artificially generated as well as in natural images especially when background noise is substantial. This superior performance can be attributed to the exclusion of false positive pixels resulting from isolated, high intensity pixels in high noise conditions. To test the method’s power in real situation we used high power confocal images of the somatosensory thalamus immunostained for Kv4.2 and Kv4.3 (A-type) voltage gated potassium channels. Moran’s method identified high intensity Kv4.2 and Kv4.3 ion channel clusters in the thalamic neuropil. Spatial distribution of these clusters displayed strong correlation with large sensory axon terminals of subcortical origin. The unique association of the special presynaptic terminals and a postsynaptic voltage gated ion channel cluster was confirmed with electron microscopy. These data demonstrate that Moran’s method is a rapid, simple image segmentation method optimal for variable and high nose conditions.

## Introduction

Differentiation of signal from noise is a key step in any image segmentation process^1^ and is based on the assumption that pixels constituting the image can be divided into spatially non-overlapping regions which are characterized by statistically similar intrinsic features^2, 3^.

During the segmentation process, the signal can be defined as a spatially congruent population of pixels with gray values, which are statistically different from the neighboring pixels, i.e., from the background^1, 3^. A theoretically optimal segmentation process should be independent of the person performing the analysis and should be based on the intrinsic properties of the image rather than manually defined values. It should consider not only the intensities but also the spatial arrangement of the picture elements (i.e., pixels).

The current segmentation methods are either based on hard or soft computing^4^. Though there is huge surge of soft computation methods based on fuzzy logic, deep learning methods neuronal networks and artificial intelligence^5–7^, image analysis based on binary, hard computation methods have their own advantages^8^ and best results can be expected from the combination of the two^9^. Many hard computing image segmentation methods depend on thresholding^4, 10^. Although more advanced threshold-based methods (TBM) utilize local not global thresholds or multilevel thresholding^11, 12^ the threshold is still defined using the intrinsic properties of the image (e.g., as deviation from the mean intensity in the picture), selection of the degree of deviation is subjective and can yield different results (Fig S1). Furthermore, even in local thresholding there is no guarantee that spatially inhomogeneous noise reaching or approaching the intensity of the signal will not be included as signal.

Here we propose a novel hard computing method for unsupervised image segmentation based on Local Moran’s spatial correlation coefficient (Local Moran’s I)^13^. The method was originally invented for geoinformatics and was used to study large-scale spatial correlation of geological^14, 15^, social or medical events e.g. suicide rates among counties ^16^ or in geomorphological studies^17^. The method, as used here, is based on the statistical evaluation of intensities among neighboring pixels, estimating the probability whether a given pixel intensity arrangement is due to a random process or not. The proposed method allows for selecting spatially congruent pixels with statistically similar intensity (i.e., signal and background) and pixels with intensity distribution not significantly different from random (i.e., noise).

In this paper, we systematically compare the performance of TBM and Local Moran’s method using images with different levels of noise. We show that, especially at high noise levels, Moran’s method convincingly outperforms TBM on both artificial and natural images. Finally, we also test Moran’s in a biological problem: clustering of voltage-gated ion channels in neuronal membranes. We show that the method can segment ion channel clusters, which are biologically relevant since their existence can be demonstrated using independent methods.

## Results

In order to demonstrate the way Local Moran’s method works for image segmentation, we used high magnification fluorescent (i.e. dark field) (Fig. 1a-j) as well as bright field (Fig 1k-t) light microscopic images showing immunoreactivity for Kv4.2 potassium channel subunit in the ventral posteromedial nucleus (VPM) of the mouse thalamus. In our approach, the unit of the analysis is the individual pixel of the image.

**Fig. 1:**
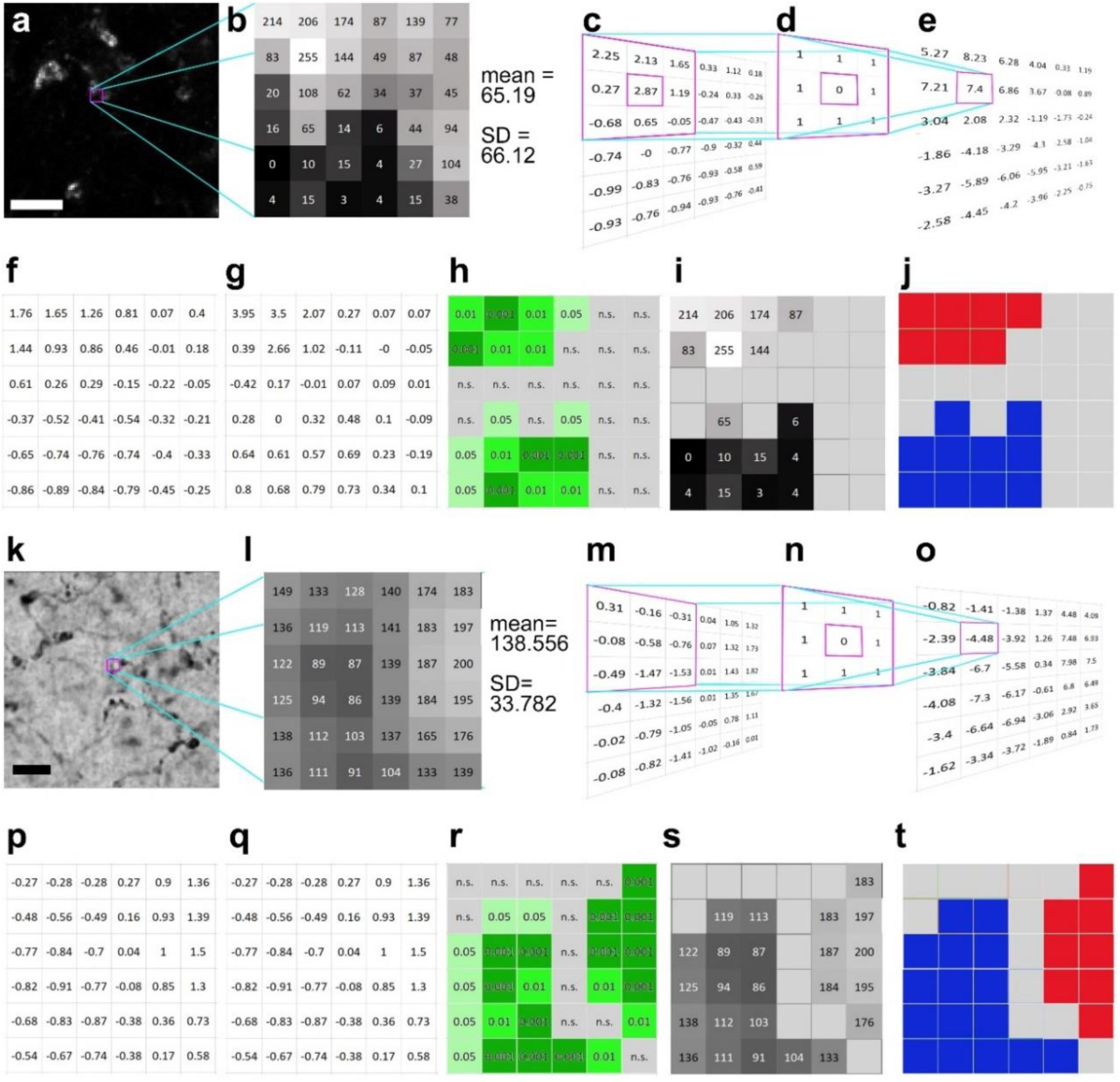
Calculation of local Moran’s I in darkfield (a-j) and brightfield (k-t) light micrographs. **a,** Large magnification fluorescent immunostaining for Kv4.2 in the VPM nucleus. **b,** A 6×6 pixel segment of the image shown in a. The squares represent the pixels with grayscale values. **c,** Normalized intensity values (z_i_). The purple framed area demonstrates the convolution step of calculations (see also methods step 1-3). **d**, A first-order convolution matrix (weight matrix, w_ij_) was used for this calculation. **e,** Weighted intensity values (w_ij_z_j_). **f,** Lagged gray values. **g,** Local Moran’s I (I_i_) values. **h,** Pseudosignificance values for each pixel. **i,** Masking the non-significant pixels (middle gray tone without numbers) resulted in two groups of pixels, where the intensity values significantly differ from a random distribution. **j**, Considering the intensity values, the upper pixel group is interpreted as an object (red), the lower one as background (blue), and the grey field between them represent an area where the pixel intensities could come from a random distribution (interpreted as noise). **k-t**, The same as a-j, but the calculations are shown on a bright field image. In this case, the low-intensity pixels representing the object are labeled with blue (t) high-intensity pixels (background) with red. Scale bars: **a**: 5 μm, **k**: 10 μm.

The method is based on testing the null hypothesis that pixel intensity distribution in the image is random, which is analyzed with the aid of convolution matrices. The results are tested by Monte Carlo simulation to identify whether the pixel intensity distribution is non-random in an image (Fig. 1). In dark field images, contiguous clustered pixels with similarly high intensities represent object (red in Fig. 1j), clustered pixels with low intensities indicate no object (e.g. lumen of a blood vessel) (blue in Fig. 1j). Group of pixels with random intensity value distribution (which do not reach significance) represent noise (grey in Fig. 1j). In bright field images, the picture is reversed, contiguous low and high-intensity pixels represent the object and no object, respectively (blue and red in Fig. 1t).

Local Moran’s I differentiates between object, no object and noise in the following way: the method first normalizes each pixel intensity value (Fig. 1c and m) (see Methods). Next, it defines a weight matrix in the local neighborhood (3×3 in this particular case) for each pixel (Fig. 1d and n), where the pixel in question receives zero weight. Then it calculates a weighted intensity value for each pixel in the image using the summed values of the neighboring pixels (Fig. 1e and o). Immediate neighbors of each pixel are assigned with the weight of 1. Next, it creates lagged grey values by dividing the matrix values with the number of neighboring pixels (Fig. 1f and p). This was called “lagged” in the original description^18^, since the distance (lag) was considered in the calculation of this particular value (which was 1 in our case). Finally, it generates the Local Moran’s I value for each pixel by multiplying the lagged gray values (Fig. 1f and p) with the normalized values (Fig. 1c and m).

The resulting Local Moran’s I value (Fig. 1g and q) will represent how the intensity of the pixel and its neighbors differs from the distribution of intensities of all pixels in the image (clusteredness). We can test the salience of this value if we repeat the same procedure with randomly selected neighboring pixels. For each pixel, the method determines the probability (pseudosignificance) of obtaining this or higher Local Moran’s I value using Monte Carlo simulations of the intensity values of all pixels in the image (Fig. 1h and r). Significant pixels (p<0.05) with high intensities (in case of dark field images) or low intensities (in case of bright filed images) will be considered as object. These steps ensures that signal is not determined by an arbitrarily defined threshold, but by the combination of spatial arrangement and pixel intensities. As a consequence, in this method consideration of each pixel as signal will not depend purely on the absolute intensity value of the pixel but on the intensity values of the neighboring pixels and the statistical probability that this spatial arrangement can arise (or not) randomly.

### Comparison of TBM and Moran’s method on artificial images

We compared the performance of Moran’s and TBM on artificial binary objects with various levels of Gaussian noise (Fig. 2a). Since in this case we defined the objects (the ground truth), True Positive Rates (TPR=true positive pixels/number of object pixels) and False Positive Rates (FPR=false positive pixels/number of background pixels) could be directly determined.

**Fig. 2:**
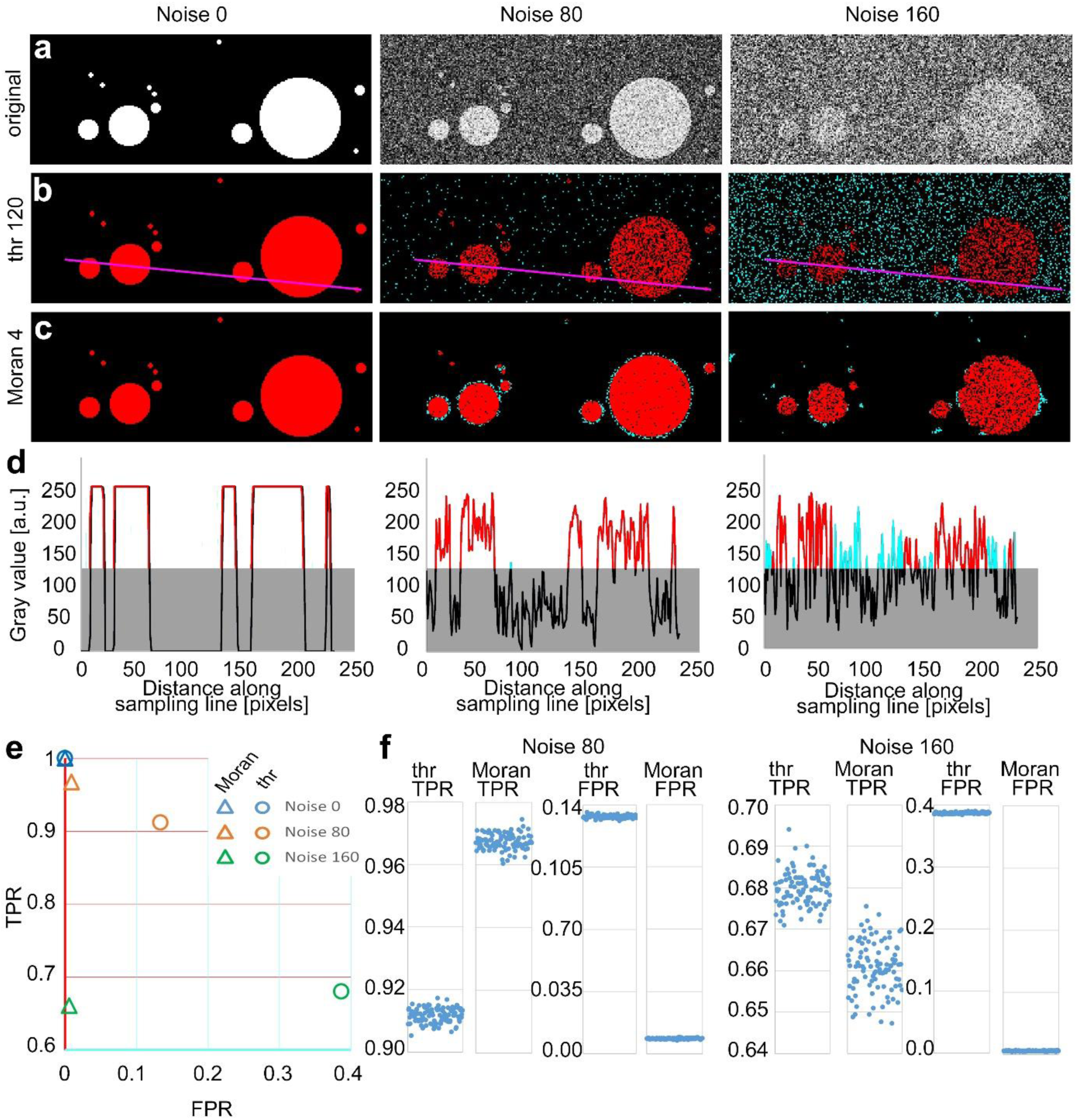
Comparison of TBM and Moran’s image selection on artificial objects. **a,** A series of artificial objects with different diameters (white circle on black background) was defined, and varying levels of noise were added (first row). **b-c,** The objects are segmented using the threshold (**b**) and Moran’s I method (**c**). Red, true positive (TP) pixels; cyan, false positive (FP); black, true negative; black true and false negative. **d,** Pixel intensity values along the sampling lines at the three noise levels. Grey rectangles indicate the threshold level. Portions of the graph’s line are color-coded (red: true positive, cyan: false positive, black: negative). **e-f,** Average TPR and FPR values of Moran’s and TBM at different noise levels from 100 random samples. Individual data points are shown separately at a different scale (**f**).

Since Moran’s method can be performed with convolution matrices of different sizes (i.e., Moran’s order, see Methods)^14^ we first determined the optimal matrix size for different object sizes at variable noise levels on circular artificial objects (Fig. S2a). First order matrix contains the immediate neighboring pixels of the central pixel (3×3), the second order contains all the immediate neighbors of the first order pixels (5×5) and so on. We found that optimal Moran’s order was independent of the noise level, and it was 0.25 times (range: 0.238-0.253) of the object size (Fig. S2b) Thus, for these measures, we selected Moran’s order 4. For the TBM, the threshold was set to 50% of the intensity level of the objects.

At zero noise, both methods had 1 TPR and 0 FPR values (Fig. 2a-c). Increasing noise levels resulted in a parallel decrease in TRP values. However, in case of the TBM FPR increased dramatically (cyan dots in Fig. 2b) due to the higher fluctuation of pixel values (Fig. 2d), which indicates a highly unreliable exclusion of false positive pixels (Fig. 2e, circles), while the FPR value in case of Moran’s method remained nearly constant (Fig. 2e triangles) with increasing noise. In case of TBM FPR increase was the consequence of that random noise frequently increased the pixel intensity values of the background above the threshold level (Fig. 2), which were detected as positive pixels.

Next, we compared the two methods in a larger sample of images (n=100) which contained 64 randomly distributed objects of different sizes. The diameter of the objects varied between 4 and 64 pixels in each image (Fig. 2a). Here, we also examined the effect of Moran’s order on the quality of detection (Fig S3). Similar to same-sized objects in images containing variable-sized objects at 0 noise level, both methods selected the objects equally well (TRP=1 SD=0, FPR= 0, SD=0, n=100 images). However, increasing the noise level resulted again in a pronounced increase in FPR in the case of TBM (Fig. 2b) whereas using Moran’s, this effect was evident only at the edges of detected objects and FPR just marginally increased even at very high noise levels (Fig. 2c). Huge increase in false positive pixels in TBM with increasing noise resulted in significantly lower FPR values compared to that of Moran’s. (Fig. 2e-f, FPR values noise 80: threshold, 0.13333±0.0009; Moran’s: 0.0085±0.0003, p<5.4*10^-215^; noise 160: threshold, 0.3866±0.0012, Moran, 0.0055±0.0005 p<2.1*10^-244^). TPR of Moran’s was also significantly better at noise 80 (Fig. 2e-f, TPR values noise 80: threshold, 0.9116±0.0023; Moran’s, 0.9674±0.0025, p<2.7*10^-130^). The TPR of TBM was marginally better at noise 160 (Fig. 2e-f, noise 160: threshold, 0.6799±0.0041; Moran’s: 0.6609±0.0060 p<5.04*10^-69^), but the difference in FPR values was so large between the two methods that it far outweighed this effect. We defined the quality of detection as a vector in the FPR/TPR space starting from the FPR=0, TPR=1 point. Quality of detection was significantly better with Moran’s method at both noise levels in a population of 100 randomly generated images (Fig. 2e FPR/TPR vector noise 80: threshold, 0.1600±2.2*10^-6^; Moran’s: 0.0337±5.8*10^-6^, p<5*10^-169^; noise 160: threshold, 0.5019±7.5*10^-6^, Moran, 0.3391±3.7*10^-5^, p<10^-156^).

Finally, to gain insight into the performance of the two detection methods in a larger parameter space, we repeated the comparison using 13 Moran’s order, 13 threshold levels and 13 noise levels (n=100 images for each condition). We found that with the exception of the lowest noise levels (0-30) Moran’s method outperformed TBM at every noise levels irrespective of the threshold mostly independent of the exact Moran’s order used (range 4-13; Fig. S4). These data demonstrate that, especially in high noise situations, Moran’s method successfully eliminates background noise and selects objects with extremely low FPR.

### Comparison of TBM and Moran’s methods on natural images

Next, we tested the performance of the two methods on greyscale images of natural objects (Fig. 3a) where, by definition, the ground truth is not known, and therefore the comparison of the two methods to a common reference is not possible. To circumvent this problem, we chose a high magnification confocal image of a nerve tissue immunostained with vesicular glutamate transporter (vGluT2), a ubiquitous marker of axon terminals with subcortical origin in the thalamus^19–21^. We prepared images with three different added noise levels and asked 23 expert human observers, who were blind to the nature of the image and the nature of the investigation, to delineate objects (Fig. 3a-b, Fig S5). Observers always started delineation in the images with the highest noise level. Next, we generated a heat map and labeled the number of experts selecting each individual pixel. For the purpose of this study, we defined ground truth as pixels which were selected by at least 12 experts (Fig. S5) and compared the performance of the two methods to this common reference (Fig. 3c-e). The size of the objects (n=36) in the image varied between 1 and 303 pixels corresponding to 0.043 and 13.003 μm^2^, respectively. For a systematic comparison of the two image segmentation methods, we analyzed the data using different threshold levels and matrix sizes (Fig. S6) at three noise levels (0, 80, 160).

**Fig. 3.**
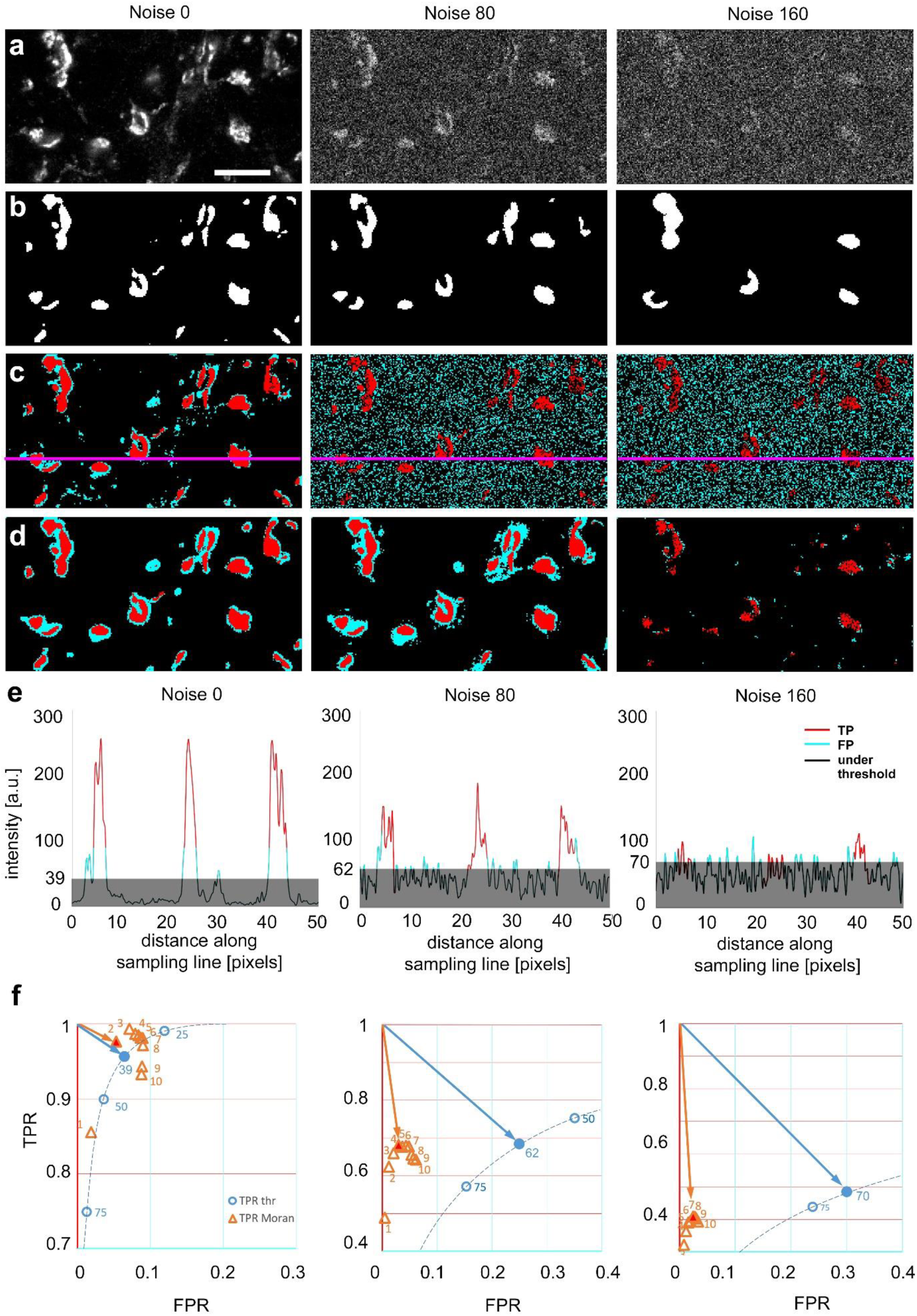
Comparison of threshold-based, and Moran’s image selection on natural objects. **a**, High-resolution confocal image (single optical plane) of Kv4.2 immunostaining in the VB complex of the mouse thalamus at three added noise levels (0, 80, 160). **b**, Majority projection of segmentation of the same images by expert human observers (n=23, see methods). Pixels selected by the majority of observers are defined as ground truth. **c,** Threshold-based segmentation with the best TPR/FPR values. **d,** Moran’s based segmentation using matrix size 4. **e,** Pixel intensity values along the sampling lines at the three noise levels. Grey areas indicate threshold levels. Portions of the graph’s line are color-coded (red: true positive, cyan: false positive, black: negative). **f,** TPR and FPR values of Moran’s and TBS at different noise levels. The threshold curve (dotted, blue line) is computed using 254 threshold levels, some of which are labeled (blue circles). Filled blue circle, optimal threshold value. Orange triangle, TPR/FPR values of Moran’s based segmentation using different matrix sizes. The size of the matrices are shown by numbers. Red filled triangle, optimal Moran’s order. Orange and blue arrows indicate the distance of the optimal points from the theoretical best point (maximal TPR, zero FPR). Scale bar a: 10 μm.

Similar to artificial objects, Moran’s method outperformed TBM at every noise level. The FPR values remained low with Moran’s method even at high noise levels at every Moran’s order tested (Fig. 3f). Thus, even though at the best threshold levels, the TPR values were comparable between the two methods, lower FPR values resulted in a better quality of detection (shorter vector) in the TPR/FPR space (Fig. 3f).

It is important to emphasize that in the case of TBM at increased noise levels, false positive pixels were homogeneously distributed in the image due to the increased background noise (Fig. 3c). In contrast, with Moran’s, most false positive pixels surrounded the human-based selection areas (Fig. 3d). This is largely the consequence of that Moran’s method detected the out of focus pixels as well, which were routinely not selected by the human experts.

These data demonstrate that when natural images were analyzed, the quality of detection was better with Moran’s method in every condition tested. The main strength of Moran’s was that it did not include pixels of random noise, like TBM.

### Application of Moran’s method for a biological problem

Next, we tested Moran’s method to address a biologically relevant problem, the clustering of Kv4.2/4.3 subunits of the voltage-gated K^+^ channels in thalamocortical neurons. Voltage-gated ion channels are either distributed homogeneously, along a gradient^22, 23^ or form high-density clusters in neuronal membranes^24–26^, endowing neurons with unique electrical properties^27–30^. However, the exact subcellular distribution of these subunits and the logic of their possible clustering in thalamic cells are unknown. Therefore, we tested whether Kv4.2/4.3 ion channel clusters, detected by Moran’s I method, form biologically relevant membrane protein aggregations.

Immunostainings for the Kv4.2 and Kv4.3 subunits displayed homogenous, diffuse labeling in the major somatosensory nucleus of the thalamus (VPM). At high magnifications, however, it became visually evident that Kv4 subunits also form strongly immunopositive clusters (2-10 µm; Fig. 4a). To study the precise subcellular distribution of Kv4 subunits, we first examined the Kv4.2 subunit at the electron microscopic level using preembedding immunogold reactions. This approach confirmed the exclusive dendritic localization of this subunit (n = 123 profiles). As suggested by the light microscopic observations, large caliber dendrites of relay cells displayed a high density of immunogold particles for the Kv4.2 subunit (n = 9 animals, Fig. 4b-c). Quantification of silver particles along the dendritic membranes (n = 3 animals) demonstrated more than three times higher density on thick (minor diameter over 1.2 µm) dendrites, preferentially innervated by large excitatory terminals containing round vesicles (RL terminals) compared to thin (minor diameter below 1.2 µm) dendrites which received very little of this type of inputs (Fig. 4d).

**Fig. 4:**
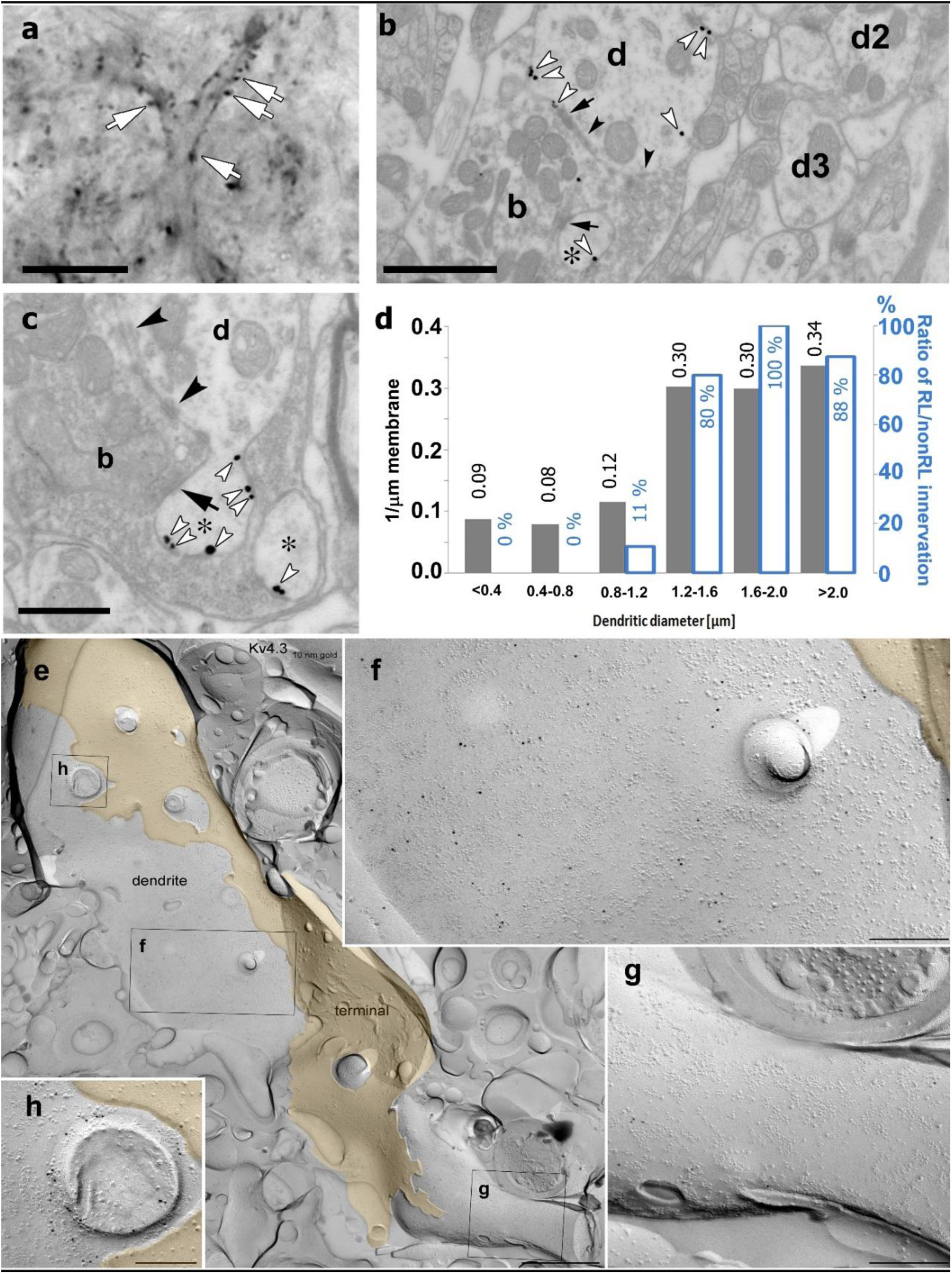
High-resolution localization of Kv4.2 subunit in the VPM nucleus of the thalamus. **a,** High-resolution light micrograph of a VPM neuron immunolabeled for the Kv4.2 subunit displaying intense immunopositive puncta (white arrows) along its proximal dendrites. **b,** An electron micrograph showing Kv4.2 immunolabeling in the VPM using the preembedding immunogold technique. A proximal dendrite, identified by its large diameter (d), has more silver intensified gold particles (white arrowheads) than thin dendrites (d2, d3). Arrows, synapses; arrowheads, puncta adhaerentia. **c,** Accumulation of gold particles in small dendritic appendages (asterisks). **d,** Quantitative analysis revealed an approximately three-fold higher density of gold particles along the membrane in large (>1.2 µm) compared to small dendrites (left y-axis, gray bars). The histogram also shows the ratio of large excitatory terminals with round vesicles (RL) innervating each dendritic category (right y-axis, white bars). **e,** Freeze-fracture replica labeling of the protoplasmic-face membrane segment of a large thalamocortical cell dendrite in VPM is shown with intense immunogold labeling for the Kv4.3 subunit. The dendrite is surrounded by a large axon terminal (yellow). Boxed areas are shown at higher magnifications in **f, g,** and **h**. **f-h,** Membrane area in the vicinity of the axon terminal contains a high density of gold particles **(f)**, whereas a neighboring area (**g**) has very few gold particles, indicating an inhomogeneous distribution of the Kv4.3 subunit. Note the intense immunogold labeling around a cross-fractured dendritic appendage **(h)**. Black arrows, synapses; black arrowheads, puncta adherentia. Scale bars: **a**: 10 μm, **b**: 1 μm, **c**: 0.5 μm, **e**, 1 µm, **f-h,** 200 nm.

To exclude the possibility that the uneven (i.e. clustered) immunogold labeling is the consequence of uneven accessibility of the antibodies to the epitopes in thick tissue and to better visualize protein aggregates at high resolution, we performed another electron microscopic experiment using the diffusion-free freeze-fracture replica immunogold labeling technique ^31^ (Fig. 4e-h). Examination of replicas of the VPM immunolabeled for the Kv4.2 or the Kv4.3 subunits revealed gold particles associated with the protoplasmic face of dendritic segments, consistent with the intracellular location of the epitopes. On large diameter dendrites, we observed uneven distribution of gold particles (Fig. 4e) consistent with our observations using confocal microscopy and pre-embedding immunogold localization. Quantitative evaluation of the reactions in one rat demonstrated that the density of gold particles for the Kv4.3 subunit was twice as high in large, compared to small diameter dendrites (12.6 ± 8.0 gold/µm^2^, n = 18 vs 6.0 ± 3.6 gold/µm^2^, n = 18). A similar difference was found for the Kv4.2 subunit, for which the density of gold particles was 22.8 ± 8.8 gold/µm^2^ (n = 9) in large diameter proximal dendrites, whereas it was 9.3 ± 3.2 gold/µm^2^ (n = 6) in small, distal dendrites. Dendritic appendages were also decorated with gold particles (Fig. 4h).

These data together demonstrate that Kv4.2 and Kv4.3 subunits display weak distal dendritic labeling, but the proximal, large diameter dendrites contain high densities of these ion channel subunits in VPM.

Next, we investigated whether the Moran’s method as described above can delineate Kv4.2 subunit clusters and quantitatively characterize them in the VPM (Fig. 5). Moran’s method clearly segmented immunopositive and negative regions (Fig. 5a-c). The immunopositive regions displayed large variability in size intensity and contained diffuse labeling and high-intensity clusters as suggested by electron microscopic investigation. To test to what extent Kv4.2 clusters detected by Moran’s method represent ion channel clusters identified on the proximal dendrites of VPM cells at the electron microscopic level (Fig. 4), we visualized large excitatory axon terminals known to be associated with the proximal dendrites of VPM cells^32^. These subcortical excitatory inputs were labeled with vesicular glutamate transporter type 2 (vGluT2, Fig. 5d)^33, 34^. We selected the proximal dendrites by performing segmentation using Moran’s method in both channels of the Kv4.2/vGluT2 double immunofluorescent images. The data showed that large, high-density clusters of Kv4.2 ion channels were indeed associated with vGluT2-immunopositive terminals (Fig.5e, cyan arrows), whereas the smaller and less intense ones were not (Fig. 5f, yellow arrows). Immunopositive clusters generally had highly variable sizes (5.59±13.26 µm^2^) and intensities (745±260 a.u.). The large majority of the non-colocalizing Kv clusters were below 1 µm^2^ and faintly stained (Fig. 5g) whereas more than the half of the colocalized Kv4.2 structures were above 5 µm^2^ and more intensely labeled (Fig. 5g). Kv4.2 clusters associated with vGluT2-positive terminals were significantly larger (12.21±18.63 µm^2^) and more intense (894±280 a.u.) than Kv4.2 regions outside the vGluT2 terminals (average size: 0.90±1.31 µm^2^, average intensity: 639±182 a.u.) indicating that Moran’s I method could delineate functionally relevant ion channel clusters associated with a specific synaptic input (Fig. 5h).

**Fig. 5:**
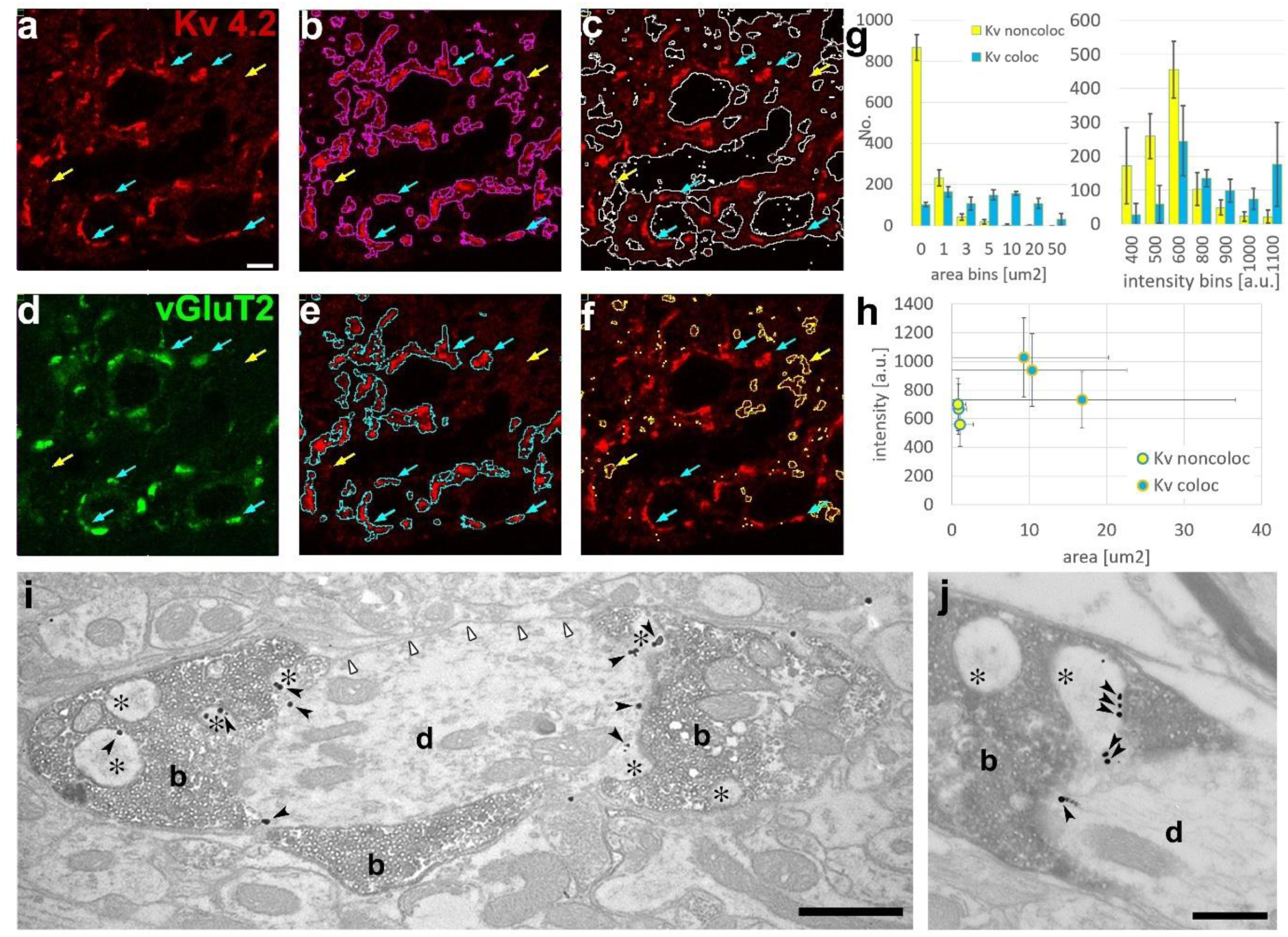
Delineation of functionally relevant Kv4.2 ion channel clusters by Moran’s method. **a,** High magnification confocal image of Kv4.2 immunostaining in the VPM of mouse thalamus. **b,** Segmentation of Kv4.2 immunopositive objects by Moran’s (magenta outlines). **c,** Segmentation of immunonegative areas by Moran’s (white outlines). **d,** Confocal image of vGluT2 immunostaining (green channel) of the same section shown in **a**. **e,** Kv4.2 ion channel clusters which overlap with vGluT2-positive terminals (cyan outlines). **f,** Kv4.2 ion channel clusters which do not overlap with vGluT2-positive terminals (yellow outlines). Blue arrows, Kv4.2 clusters overlapping with vGluT2 terminals; yellow arrows, Kv4.2 clusters non-overlapping with vGluT2 terminals (holds for a-f). **g,** Histograms of the colocalizing and non-colacalizing Kv 4.2 immunopositive boutons pooled from three animals segmented by Moran’s method and binned by area (left) and intensity (right). Yellow bars, Kv4.2 objects non-overlapping with vGluT2; cyan bars, Kv4.2 objects overlapping with vGluT2. **h,** Scatterplot of area vs staining intensity from three different animals. Yellow dots, Kv4.2 objects non-overlapping with vGluT2; cyan dots, Kv4.2 objects overlapping with vGluT2. **i, j** Silver particles (black arrowheads) indicating Kv4.2 immunoreactivity are localized on dendritic appendages (asterisks) and dendritic shafts (d) contacted by vGluT2-positive terminals (b, DAB positive) in the VPM. The plasma membrane of the same dendrites that are not contacted by vGluT2 positive terminals (white arrowheads) contains very few particles for the Kv4.2 subunit. Scale bars: a-f: 5 μm **i**: 1 μm, **j:** 0.5 μm

To verify the association of afferents and ion channel clusters, we performed double immunostaining for Kv4.2 and vGluT2 at the electron microscopic level (Fig. 5i-j). Indeed, membrane-associated silver intensified gold particles for the Kv4.2 subunit were associated with membrane surfaces covered by vGluT2 positive terminals, as demonstrated by the Moran’s method at the light microscopic level. Membrane segments contacted by vGluT2 boutons showed more than 4 times higher gold particle density (37.0 particles/100 μm membrane profile, total measured length: 601 μm) compared to surrounding membrane surfaces that were not contacted by vGluT2 positive axons (8.9 particles/100 μm, total measured length: 394 μm). These data reveal a novel association of voltage-gated ion channel clusters on the postsynaptic membrane surfaces with a specialized presynaptic terminal.

## Discussion

In this study, we demonstrate that Local Moran’s method is suitable for image segmentation and outperforms TBM in both artificial and natural images, especially when the noise is substantial. We used this method to successfully delineate and quantify biologically relevant membrane protein clusters. Since Moran’s selections are based on the relative intensity values of neighboring pixels, it can select contiguous points as objects, unlike TBM. TBM does not use spatial information of the pixels. Therefore, it selects random noise as well, provided it reaches the threshold. Unlike in TBM, selection of the signal will not depend on the absolute intensity of its pixels in Moran’s method since the image processing starts with a normalization step. This way, it uses inherent properties of the image to select clusters. This will allow the systematic, quantitative comparison of images where sample preparation and image acquisition cannot be standardized, such as in most histological samples.

Local Moran’s method has not been used for image segmentation so far. Its global version, however, has already been utilized in a wide range of biomedical image analyses. Global Moran’s method assigns a value to the whole image and describes whether pixel distribution is random or not, but does not segment pixels to object, no object and noise. Global Moran’s I were used to examine muscle and fat tissue on 3D MRI images^35^ or to measure anisotropy in ultrasound imaging^36^. In addition, Global Moran’s method was also used to analyze the homogeneity of the temporal information in Ca^2+^- signals in periodontal fibroblasts^37^ or to detect correlated neuronal activity^38^. Based on these, the utilization of Local Moran’s method, as described here, will not be restricted to segmenting objects on 2D images but can be extended for 3D image segmentation and analyzing temporal signals.

The strongest argument to use Local Moran’s I instead of threshold-based segmentation is the effective elimination of false positive pixels from the background. Although the intensity of noisy pixels can reach or exceed that of the objects, white noise does not cause a pattern. Detection of the lack of spatial correlation is a powerful tool to ignore false positive salient pixels in the background.

As indicated above, Moran’s method works without user-dependent intensity values. However, the size of the convolution matrix needs to be set by the user. In this study, we examined the effect of matrix size in a wide range of applications (Fig. 3f, Fig. S2-4). We determined that selection of the signal is relatively insensitive of the exact matrix size within a certain range (Moran’s order 3-8), especially in noisy conditions. A priori knowledge of the expected object size helps to define the best matrix size since we found a linear correlation between object size and Moran’s order for optimal image segmentation (Fig. S2b).

We found that Local Moran’s method can delineate and quantitatively characterize Kv4.2 ion channel clusters in the somatosensory thalamus formed in the proximity of large vGluT2 positive afferent nerve terminals. The selective association of a given presynaptic terminal type and the postsynaptic accumulation of ion channel clusters was verified by independent electron microscopic methods. The functional relevance of this unique association remains to be determined: it may participate in timing the precision of sensory integration, but recently the non-synaptic role of voltage-gated ion channel accumulation has also been proposed^39^.

## Methods

### Animals

Six adult male C57Bl6 mice were perfused to examine the Kv4.2 expression. All animals were bred and kept in the Animal Facility of the Institute of Experimental Medicine, Budapest, Hungary. All procedures with mice were approved by the Animal Welfare Committee of the Institute of Experimental Medicine, Budapest, conformed to guidelines established by the European Community’s Council Directive of November 24, 1986 (86/609/EEC). Experiments were authorized by the National Animal Research Authorities of Hungary (PE/EA/877-7/2020).

The animals were deeply anesthetized with a mixture of ketamin, xylazine (Produlab Pharma BV, Raamsdonksveer, The Netherlands) and promethazine (Pipolphen, EGIS, Budapest, Hungary; 2 mg/kg, 10 mg/kg and 5 mg/kg body weight, respectively) and perfused through the heart using a fixative containing 2% or 4% paraformaldehyde (TAAB, Berkshire, UK) and 15% saturated picric acid (Spektrum-3D Kft, Debrecen, Hungary) dissolved in 0.1 M phosphate buffer (PB).

### Immunohistochemistry, light microscopy

Coronal sections (50 µm) were cut from the thalamus, and thoroughly rinsed in 0.1 M PB followed by 0.1 M tris-buffered saline (TBS, pH=7.4). For the investigation of the distribution of A-type K^+^ channel subunits in different nuclei of the thalamus, the following antibodies were used; mouse anti-Kv4.2 (from NeuroMab Inc, UC Davis, CA; 1:300, code 75-016 and 75-017); rabbit anti-Kv4.2 (from Alomone labs, Jerusalem, Israel, 1:500, code APC-023 and APC-017). This was followed by the incubation with biotinylated horse anti-mouse or goat anti-rabbit antibodies (bHAM or bGAR; Vector Laboratories, 1:300; in TBS; 3 hours, code BA-2000 and BA-1000, respectively) and by avidin-biotinylated horseradish peroxidase complex (ABC, Vector, 1:300; 90 min, code PK-6200). The immunoreactions were visualized using diaminobenzidine (DAB, Sigma) or nickel intensified DAB (DAB-Ni) as chromogens. Sections were treated with OsO_4_ (1% in 0.1 M PB; 40 minutes, EMS, Munich, Germany), dehydrated in ethanol and propylene oxide, and embedded in Durcupan (ACM, Fluka, Buchs, Switzerland).

For the visualization of the vesicular glutamate transporter 2 (vGlut2) and the Kv4.2 subunit, fluorescent double labeling experiments were carried out. The sections were incubated with a mixture of mouse anti-Kv4.2 (1:300) and rabbit anti-vGlut2 (1:1500; Synaptic Systems, Göttingen, Germany, code 135 402) or rabbit anti-Kv4.2 and mouse anti-vGlut2 (1:3000, Millipore, Temecula, CA, code MAB 5504). This was followed by a mixture of Alexa488 conjugated goat anti-rabbit (A488-GAR; 1:500; 2.5h; Molecular Probes, Leiden, The Netherlands, code A-11001) and Alexa594 conjugated goat anti-mouse (A594-GAM; 1:500; 2.5h; Molecular Probes, Leiden, The Netherlands, code A-11012) IgGs. In some experiments Cy3 conjugated goat anti-mouse (1:500 Jackson ImmunoResearch Europe Ltd., Suffolk UK, code 111-165-144) IgGs were used instead of Alexa594. After the reactions, the sections were mounted on glass slides in Vectashield (Vector Laboratories, Burlingame, CA) and were examined using a Zeiss Axioplan2 microscope (wavelengths for filter set absorption and emission in nm: 365 bandpass/420–460; 450–490/512–542; 540–546/578–643). Digital micrographs were taken with a digital camera (Olympus, DP70, Tokyo, Japan). Z image-stacks were also acquired with a confocal laser scanning microscope (Olympus FV1000 or Nikon ECLIPSE Ni-E).

### Freeze-fracture replica labeling

80 μm coronal sections were cut from the forebrain of a perfusion fixed Wistar rat (2% paraformaldehyde and 15 v/v% picric acid in 0.1 M BP) and small blocks from the VPM were cut out for rapid freezing with a high pressure freezing machine (HPM100, Leica Microsystems) and were fractured and coated as described earlier^23^.

Replicas were digested in a solution containing 2.5% sodium dodecil sulphate (SDS) and 20% sucrose in TBS (pH = 8.3) at 80°C for 18 hours. Following several washes in TBS containing 0.05% bovine serum albumin (BSA), replicas were blocked in TBS containing 5% BSA for 1 h, then incubated overnight in blocking solution containing either a rabbit anti-Kv4.3 (1:500; Chemicon/Millipor, code AB5194) or a rabbit anti-Kv4.2 IgG (1:500; Alomone Labs, code APC-023). After this, the reactions were visualized with goat anti-rabbit IgGs coupled to 10 nm gold particles (1:100; British Biocell International).

### Calculation and properties of Moran’s I

The code for the analysis is available online as a user-friendly MATLAB script at: https://github.com/dcsabaCD225/Moran_Matlab/blob/main/moran_local.m

In details, methods for detecting spatial inhomogeneities (clustering) in geostatistics were described long ago^14, 18^ and they were adapted biomedical research much later. There are only sporadic papers using Moran’s spatial correlation coefficient (Global Moran’s I) for biomedical image analysis in order to detect clustering in general^40^ without separating particular clusters. We were interested in localization of clusters that is also possible with the help of a special case of local indicators of spatial association named Local Moran’s I ^18^. It can be calculated for each individual pixel of the image:

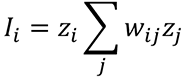

where z_i_ is the observed pixel intensity value, z_j_ are the neighboring pixel intensity values weighted by w_ij_. The minimal size of detectable particles is heavily influenced by the size of the weight matrix^41^. In our case we considered the immediate neighboring pixels (j) of each pixel when calculated (i) with an equal weight of 1 in first order matrices. In case of larger matrices, the weights were assigned according to a Gaussian distribution. This coefficient (I) gives an estimate of the homogeneity of the spatial distribution of pixels with similar gray values. Morans’s I is close to zero if the distribution is random, whereas it has a value around −1 if the distribution is uniform and scattered, and +1 if the distribution is uniform and clustered. The result can be statistically tested by a permutation test, where the data points (pixels) are randomly redistributed and the same coefficient is calculated for each permutation. Then a pseudo significance is computed as p=(G+1)/(R+1), where G is the number of instances where the absolute value of the observed I is higher than or equal to the absolute value of the original I, and R the number of repeats. For example, if none of 99 permutations have a higher I value, then the pseudo p-value is 1/100=0.01.

Based on the principles above we performed the following step-by-step procedures of Moran’s analysis.

1. Normalization of pixel intensity values (V) z_i_=(V-Vm)/SD where z_i_ is the normalized intensity value, V is the original intensity value, Vm is the mean intensity of all pixels and SD is the standard deviation of intensity of all pixels (Fig. 1a-c)
2. Creating weights (w_ij_) For each pixel, a set of neighboring pixels was defined. These pixels got the weight 1, all others including the actual central pixel got the weight 0. For the present work, we considered the immediate 8 neighbors of the pixels, in other words we applied a first order queen contiguity. (Fig. 1d)
3. Calculating weighted intensity values (w_ij_z_j_) After multiplying the weights (w_ij_) with the appropriate normalized pixel intensities (z_j_), each weighted intensity values belonging to a central pixel were summed (Σ_*j*_*w_ij_z_j_*) (Fig. 1e)
4. Calculating the lagged gray values that are used for the scatter plots lgv=Σ_*j*_*w_ij_z_j_*/n where n is the number of neighbors considered (in the corner n=3, on the edge n=5, else n=8, Fig. 1f).
5. Multiply the lagged gray values with the normalized intensity values gives the Moran’s coefficient (I_i_) for each pixel. (Fig. 1g)
6. Calculation of pseudo-significance (p) The pixels of the image were randomly distributed except the central one and the Local Moran’s I is calculated again. The pseudo significance level (see above) was calculated for each pixel. The pixels where the p is extreme low, considered to be surrounded by similar pixel values, thus members of homogenous, non-random pixel groups (Fig. 1h).
7. Defining clusters The non-random pixel groups can be divided in further sub-groups according to the relation of the pixel intensities of their central (z_i_) and neighboring (lagged gray value) pixels: high-high, high-low, low-high and low-low. If both values are high, then this group is regarded a cluster (Fig. 1i-j).

Previously this method was used for analyzing geographical distribution of medical data only, but univariate Local Moran’s I seems to be suitable for delineating clusters of pixels with similar gray values (either high or low) as outliers from the assumed random distribution. The results can be mapped as a graded image and can be used for observer-independent identification of fluorescent clusters. We selected the biologically meaningful spots, which might correspond to pre-or postsynaptic ion channel clusters if they meet the size criteria defined by electron microscopic observations.

### Matrix Size

The convolution matrix size defines the number neighboring pixels considered in the convolution step. Traditionally the size of the matrix is described as the order of the Moran’s test. The first order matrix contains 1 pixel in all directions of a central pixel, i.e. size of the matrix is 3×3, the second order matrix contains the first order pixels and their neighbors, therefore the size is 5×5, etc (Fig S6, orange transparent squares). The matrix size has reportedly a strong effect on Global Moran’s I results ^41^.

Thus, we determine optimal matrix order in our Local Moran’s calculation, to find out which provided the best object selection under different noise levels. Each object size (2, 4, 8, 16, 32, 64 pixels in diameter) was tested with 13 different Moran’s order (1-10, 16, 32, 64) (Fig. S3a). The data indicated that independent of noise levels, the best Moran’s order is 0.25 times the object size (Fig. S2b).

### Manual delineation

Twenty-three volunteers were asked to delineate objects on images. The volunteers did not have any a priori knowledge about the images. The only information that they had is it contains light objects on dark background. Each volunteer received three images in the same order. All three images were the same micrograph of vGluT2 immunostained boutons but with different level of noise (SD of noise was set 160, 80, 0, in this order). Each volunteer must have completed the delineation of all objects of the noisiest image (Fig. S5a) first, before receiving the less noisy and finally the noiseless image (Fig. S5b-c). Human observers drew quite different borders (Fig. S5g-i), but the majority projections (set of pixels selected by the majority of participants, i.e. at least 12 observers) of these object masks resembled to the original structure even in case of the noisiest image (Fig. S5d), but of course it was much clearer in case of the noiseless image (Fig. S5f). Since the biological samples does not have a similar ground truth like the generated objects, we used the areas of such projections of human delineations as a control in the further tests.

## Acknowledgements

This research was supported by the Wellcome Trust (ZN, LA). In addition, LA was supported by an ERC Advanced Grant (FRONTHAL, 742595) and the European Union project RRF-2.3.1-21-2022-00004 within the framework of the Artificial Intelligence National Laboratory and Lendület_2023_90. ZN is the recipient of a Hungarian Academy of Sciences Momentum Grant (Lendület, LP2012-29) and an ERC Advanced Grant (293681). We thank the Light Microscopy Center at Institute of Experimental Medicine for kindly providing microscopy support. Authors would like to express their deepest gratitude to Prof Luc Anselin (Center for Spatial Data Science, University of Chicago) and Dr Szabolcs Káli (Instiute of Experimental Medicine, Budapest) for the valuable discussion about analysis of spatial association, and to Krisztina Faddi for the excellent technical assistance.

---

**Fig. S1.**
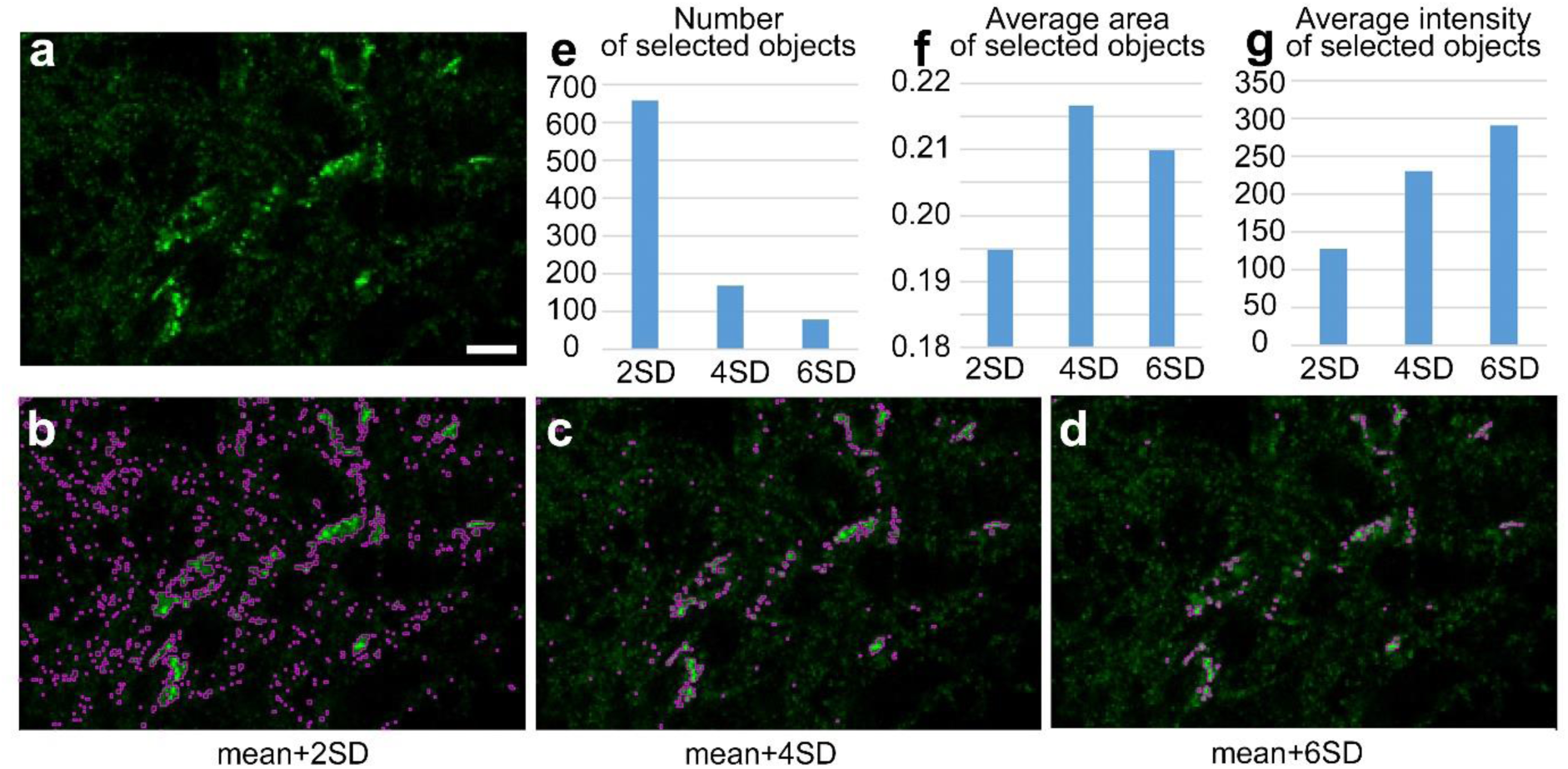
Effect of threshold level on object detection. **a,** High power confocal image of Kv4.2 immunostaining in the ventrobasal complex of the mouse thalamus. **b-d,** Segmented objects (purple) of the same picture at three different threshold levels. **e-g,** Number of objects (**e**, No), average area of selected objects (**f**) and average pixel intensity values of the selected objects (**g**). Note the large difference in every measure depending on the threshold level. Scale bar a: 5 μm.

**Fig. S2.**
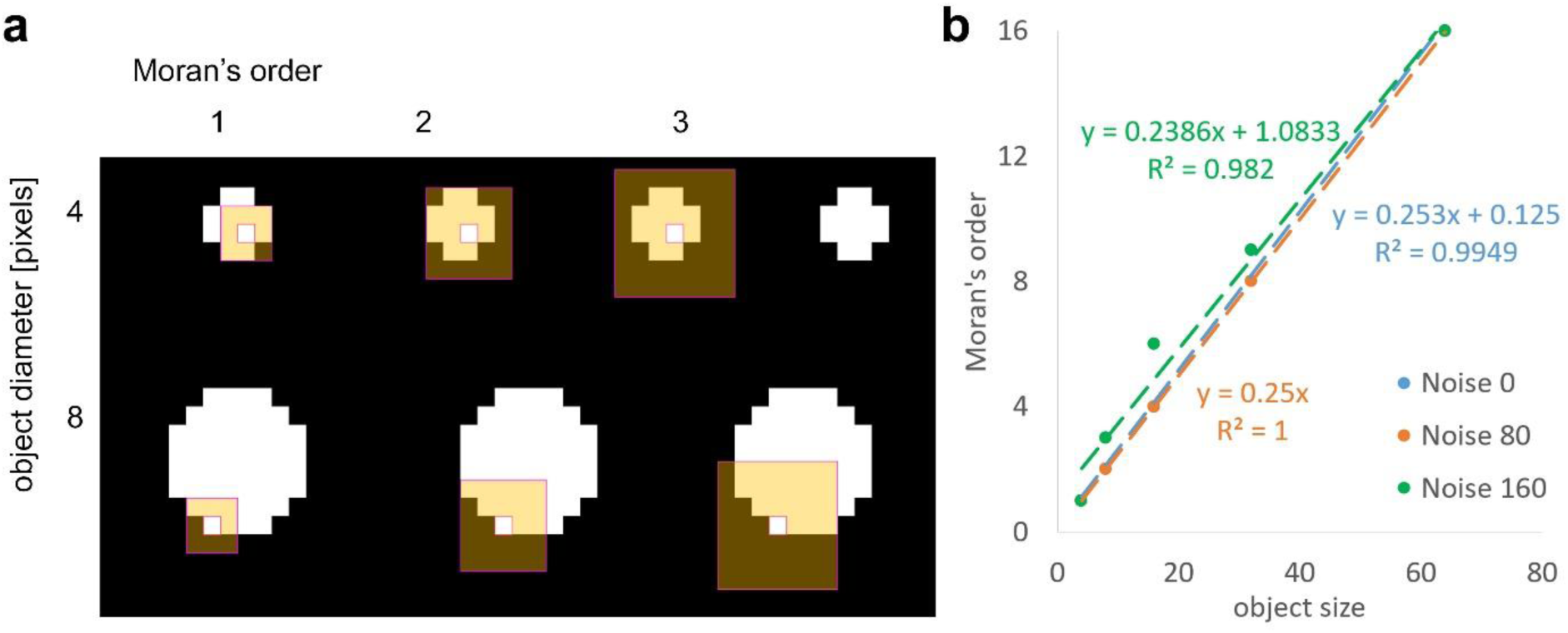
Defining optimal Moran’s matrix. **a,** Sizes of convolution matrices with different order (1, 2, 3; orange squares around the central pixel) with artificial objects (white) of different sizes (4 and 8 pixels in diameter) used in Moran’s tests. **b,** Optimal order number in relation to the object size (4-64) at different noise levels. To determine the optimal Moran’s order, we examined the quality of detection of objects with variable object sizes (diameter 4-64 pixels) at different noise levels. The optimal order was plotted against the object size. The points were fitted on a line with a slope of approximately 0.25, independent of the noise level.

**Fig. S3.**
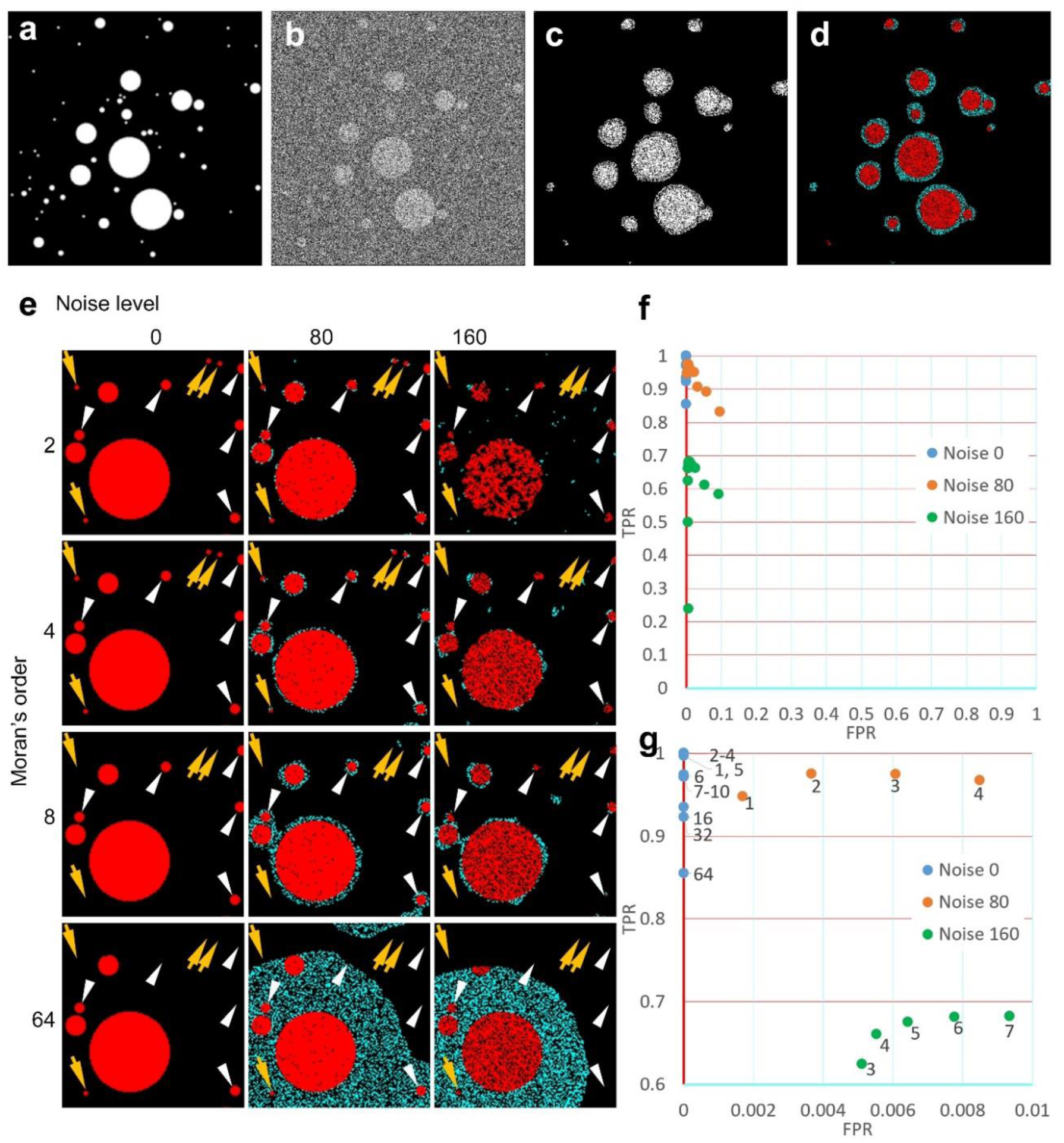
The effect of Moran’s order on the quality of detection using variable object sizes and different noise levels. **a,** One image out of the 100 containing randomly placed artificial objects with diameters ranging from 4-64 pixels. **b,** The same image with added noise. **c,** Objects selected by Moran’s method with a 16^th^ order convolution matrix. **d,** The same selection pseudocolored with red (true positive) and cyan (false positive). **e,** Similar objects were investigated at different noise levels and with different orders of Moran’s test. Note the increase of false positive rates at high noise levels and higher Moran’s order. Yellow and white arrows indicate small objects that became undetected at higher Moran’s order. **f,** Effect of matrix size on the quality of detection (in a TPR/FPR space) in images containing 100 objects with variable size at three different noise levels. **g,** Zoomed in version of the plot shown in **f**. Numbers indicate the matrix size.

**Fig. S4.**
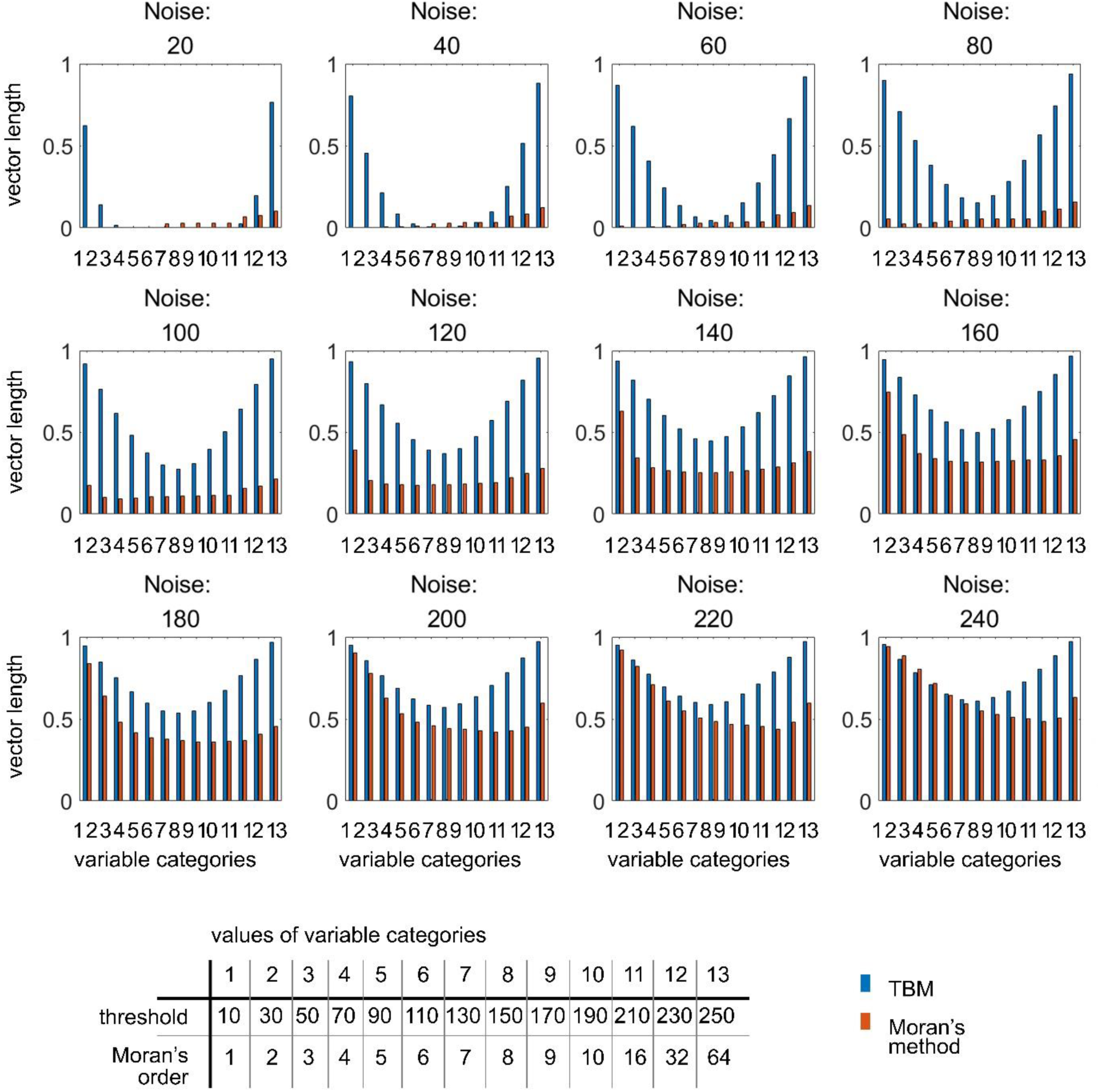
Comparison of the vector length at various noise levels (20-240) from the theoretical best (0, 1 point in the TPR/FPR space) to the closest point of the TPR/FPR curve in case of the TBS and Moran’s methods using images containing randomly placed artificial objects with diameter ranging from 4-64 pixels. Blue: TBM, red: Moran’s method. The bins of the x-axis (1-13) indicate different x-values for the two methods (see the table at the bottom) for comparisons. Threshold levels vary between 10-250 for TBS (blue bars), Moran’s orders between 1-64 (orange bars). Shorter vector length indicates better performance. Note that between 60 and 160 noise levels the exact value of Moran’s order does not affect the quality of detection and all Moran’s values are below the TBS values.

**Fig. S5.**
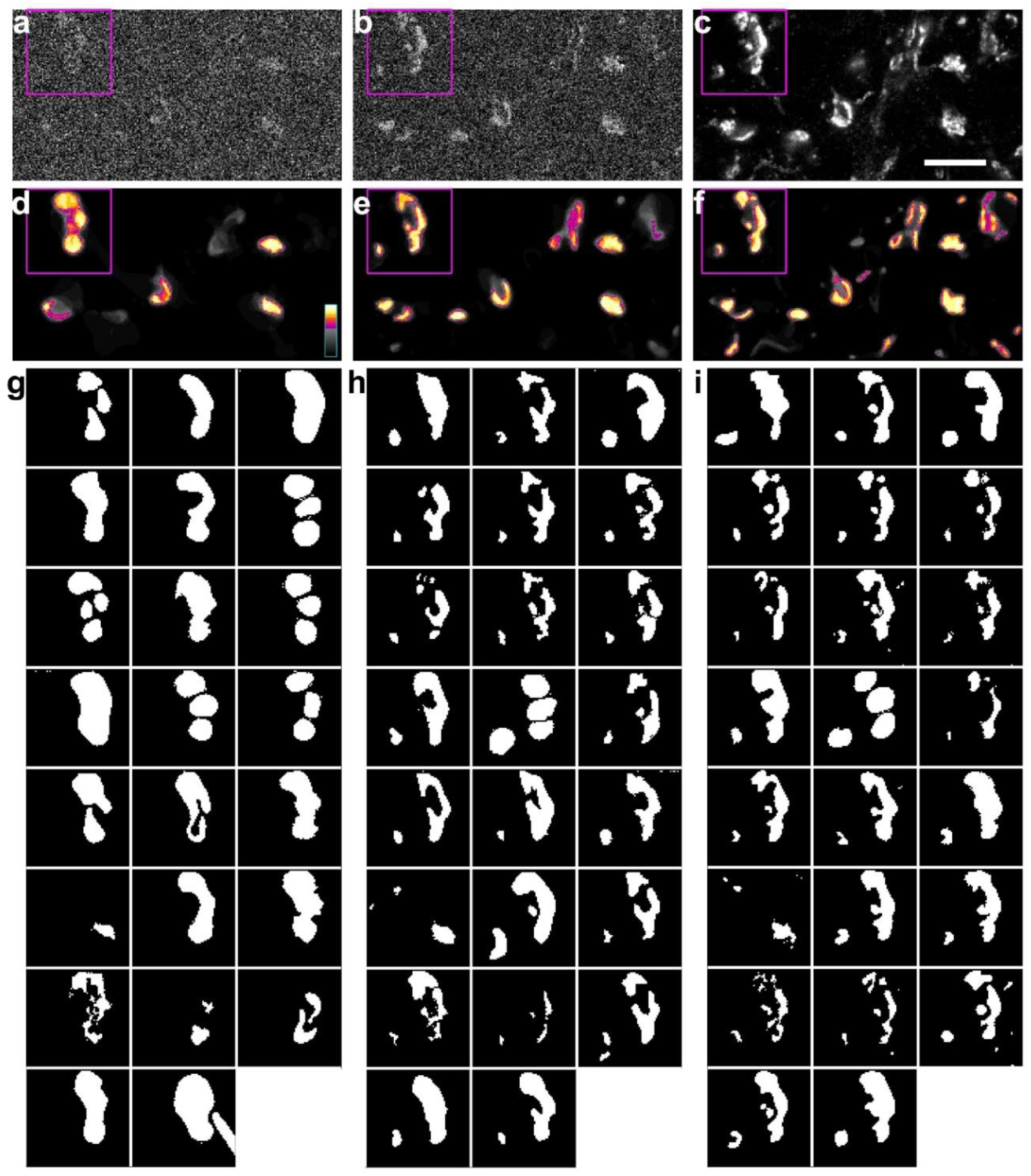
Definition of ground truth on natural images. **a**-**c**) Micrograph of fluorescently labeled Kv4.2 ion channel clusters with added noise (**a**, **b**, **c**, noise level 160, 80, 0, respectively) in the order in which they were presented to the human observes. **d**-**f**) Selection of objects by 23 observers who were blind to the nature in the same images. Gray pixels were selected by less than half of the participants (n=23), the pseudocolored pixels were selected by more than half and were referred as “ground truth” in this study. **g**-**j**) Selection of objects within the framed areas (magenta rectangles in **a**-**c**) by each of the 23 human observer. Each observer outlined first the noisiest image (**g**, rectangle in **a**), then the less noisy one (**h**, rectangle in **b**), finally the original input image (**I**, rectangle in **c**). Scale bar c: 10 μm.

**Fig. S6.**
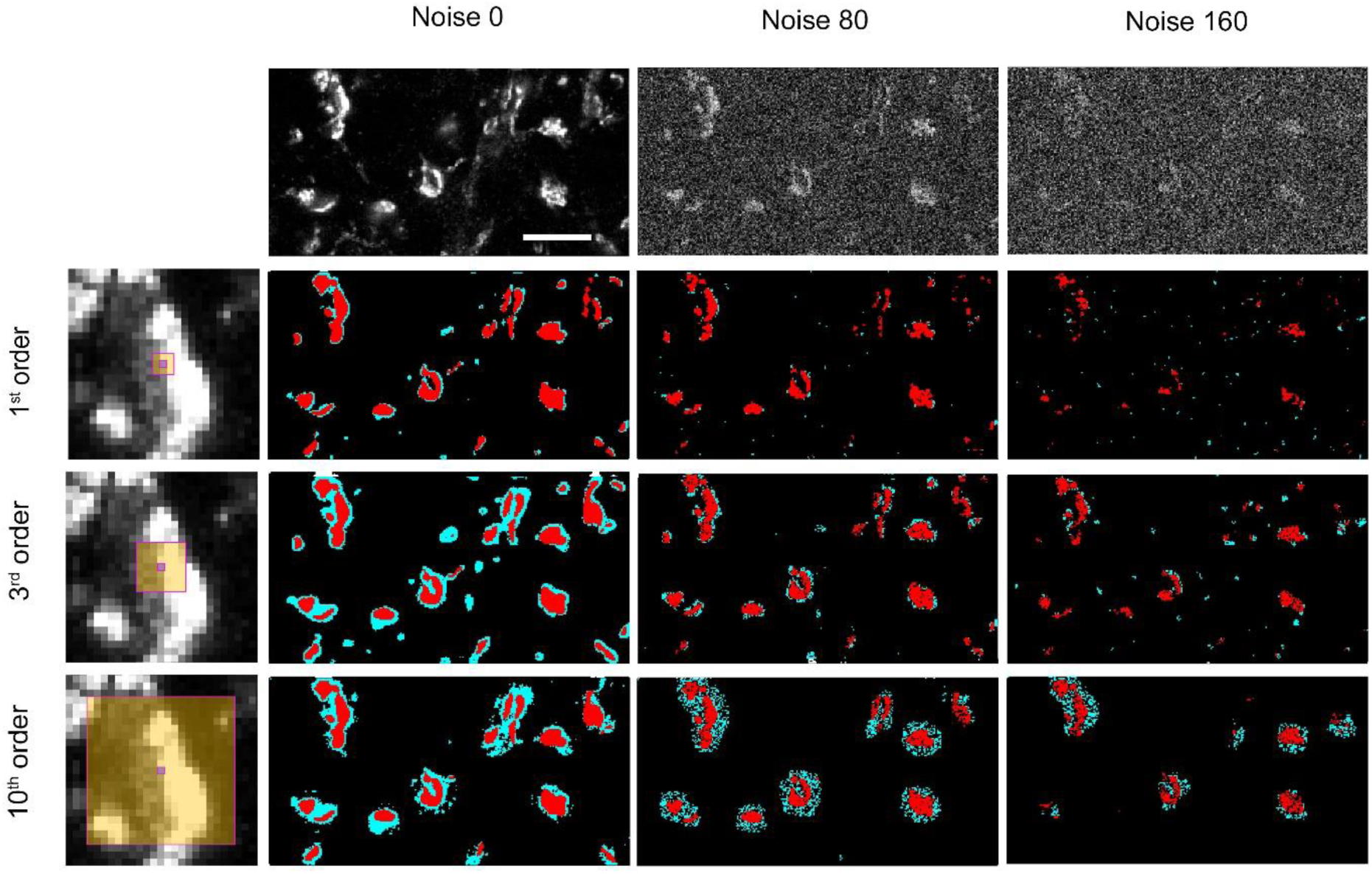
Effect of convolution matrix size on the visual results. Natural image (Kv4.2 immunoreactive boutons) at three different noise levels (0, 80, 160). The putative objects were detected with different convolution matrices (1^st^, 3^rd^, 10^th^ order). The sizes of matrices are displayed on one of the boutons for the sake of comparison. Red: true positive pixels, cyan: false positive pixels. Scalebar 10 μm.

